# Intermediate antiparallel beta structure in amyloid plaques revealed by infrared spectroscopic imaging

**DOI:** 10.1101/2023.04.18.537414

**Authors:** Brooke Holcombe, Abigail Foes, Siddhartha Banerjee, Kevin Yeh, Shih-Hsiu J. Wang, Rohit Bhargava, Ayanjeet Ghosh

## Abstract

Aggregation of amyloid beta (Aβ) peptides into extracellular plaques is a hallmark of the molecular pathology of Alzheimer’s disease (AD). Amyloid aggregates have been extensively studied in-vitro, and it is well known that mature amyloid fibrils contain an ordered parallel β structure. The structural evolution from unaggregated peptide to fibrils can be mediated through intermediate structures that deviate significantly from mature fibrils, such as antiparallel β-sheets. However, it is currently unknown if these intermediate structures exist in plaques, which limits the translation of findings from in-vitro structural characterizations of amyloid aggregates to AD. This arises from the inability to extend common structural biology techniques to ex-vivo tissue measurements. Here we report the use of infrared (IR) imaging, wherein we can spatially localize plaques and probe their protein structural distributions with the molecular sensitivity of IR spectroscopy. Analyzing individual plaques in AD tissues, we demonstrate that fibrillar amyloid plaques exhibit antiparallel β-sheet signatures, thus providing a direct connection between in-vitro structures and amyloid aggregates in AD brain. We further validate results with IR imaging of in-vitro aggregates and show that antiparallel β-sheet structure is a distinct structural facet of amyloid fibrils.

## Introduction

Alzheimer’s disease (AD) is a rapidly growing public health challenge worldwide. Nearly six million Americans have Alzheimer’s dementia, and the cost of care is estimated to be more than a quarter of a trillion dollars^1^. Currently no cure for AD exists; therapeutic interventions and care are mostly palliative in nature^2^. Aggregation of the amyloid beta (Aβ) protein into extracellular plaques is one of the main pathological features of AD^3-5^. Amyloid plaques can be broadly divided into dense-cored and diffuse morphologies. It is generally believed that cored plaques contain fibrillar amyloid aggregates, while amorphous prefibrillar aggregates constitute diffuse plaques^3, 4, 6^. The structure of Aβ aggregates formed in-vitro has been the subject of many detailed investigations, which have conclusively identified that mature Aβ fibrils have parallel β-sheet structure while prefibrillar oligomers may have antiparallel β-sheet structure ^7, 8-11,^. However, deviations from these structures have also been demonstrated. The Iowa mutant of Aβ, which is implicated in Cerebral Amyloid Angiopathy (CAA), wherein amyloid aggregates accumulate in blood vessels, has been shown to form transient antiparallel fibrils^12^. Recent studies using nanoscale infrared (IR) spectroscopy have identified the presence of antiparallel structure in early-stage fibrils and prefibrillar aggregates of wild-type Aβ^13-16^. Certain brain-derived Aβ fibrils, formed from seeded growth using AD tissue extractions, have also been found to contain antiparallel β-sheet arrangement^17, 18^. However, it is yet to be fully understood how translatable these findings are to different amyloid structures in plaques in AD brain tissues. Antiparallel fibrils have been shown to be neurotoxic compared to their parallel counterparts^19^; hence, it is important to understand if and how these structural deviations manifest themselves in AD plaques and correlate with onset and variation of neurological symptoms^20^. However, detailed information about Aβ aggregates in AD brain is lacking and key questions, specifically regarding their structural distributions, remain unanswered. While it can be expected that amyloid aggregates in plaques can exist in a range of structures as observed in vitro, it is currently not known if this is true or if the same amyloid structure and chemical composition persist across plaques for a given patient. It is also not known how this structural distribution correlates with onset and different levels neurological symptoms between different patients. Understanding the protein structure in plaques in AD brain tissues represents an important and significant problem which can enable the development of targeted therapeutics. The major impediment towards probing protein structures directly in ex-vivo tissue specimens has been the lack of spatial resolution of common structural biology techniques, without which, plaque-specific observations cannot be made. IR spectroscopic imaging, which allows for spatially-resolved molecular measurements through their absorption spectra, is ideally suited to circumvent this limitation. IR imaging strategies have been employed to investigate plaques in human and mice AD brain tissues^21, 22-25^; however, most of these reports have been limited to proof-of-concept studies. Additionally, the applicability of findings from mouse models to the course of disease in humans is debated. Recent advances in IR imaging technology have been used to study larger sample sizes of amyloid plaques in human AD brain^23-25^ and has shown that fibrillar plaques indeed contain parallel β structure. Furthermore, structural variations can also exist between different amyloid aggregates in the AD brain. However, the presence of toxic intermediates observed in-vitro has never been specifically probed in plaques. In this report, we address this gap in knowledge and demonstrate that antiparallel transient intermediates can exist in amyloid plaques, thus providing the first direct link between in-vitro and AD tissue plaque structures.

## Results and Discussion

For this study, we investigated the IR spectral maps of 80 plaques from the frontal lobe of two AD human tissue sections. Plaques on the tissue sections were identified using immunohistochemical (IHC) staining with an anti-amyloid β antibody (MOAB-2)^26, 27^. Amyloid plaques can be categorized broadly into cored and diffuse morphologies based on the histological stained images. It is well known that cored plaques contain fibrillar aggregates of Aβ at their core, which is the most dense, central portion of the positively stained region^4, 6^. In this work, we specifically focused on cored plaques to identify variations in secondary structures of fibrillar aggregates in AD tissues. Figure 1A shows the IHC stained image of several representative cored plaques. Representative point spectra from the plaque cores of the Amide-I vibrational mode, which is known to be reflective of protein secondary structure^28-30^, are plotted in Figure 1B. The spatial locations from where the spectra are extracted are marked in Figure 1A. The spectra clearly indicate that there are significant plaque-to-plaque variations that differ from non-plaque tissue (shown in black). The non-plaque tissue exhibits a peak at ∼1660 cm^-1^, typically seen in IR spectroscopy of fixed tissues^24, 31, 32^, and is usually attributed to a mixture of unordered random coils and β-turn secondary structures. We also observe a shoulder at ∼1630 cm^-1^. The plaque marked in blue exhibits an IR spectrum identical to non-plaque tissue, whereas the spectra of the green and red marked plaques exhibit increased intensity at ∼1630 cm^-1^. In addition, the red spectrum also contains a distinct shoulder at ∼1692 cm^-1^. It is well known that β-sheet secondary structures, which are the main constituent of amyloid fibrils, have a characteristic absorption at ∼1630 cm^-128-30^. Therefore, the observation of enhanced β-sheet signatures from plaque cores is consistent with the view that plaque cores contain fibrils. Additionally, a peak or shoulder at ∼1692 cm^-1^ is an established signature of protein secondary structure and can be attributed specifically to antiparallel β-sheets^15, 28, 33^, whereas the band at ∼1630 cm^-1^ can arise from both parallel and antiparallel β-sheets and is reflective of overall β-sheet structure contribution to the spectrum. Thus, a plaque exhibiting enhanced absorbance at ∼1630 cm^-1^ suggests the presence of β-sheets, either parallel or antiparallel, while enhanced absorbance at both ∼1630 cm^-1^ and ∼1692 cm^-1^ specifically points to antiparallel β-sheet structure. In-vitro measurements have conclusively determined the secondary structure of amyloid fibrils as parallel β-sheets^8, 10, 11, 33-35^; thus, finding antiparallel β-sheet signatures in plaques was not expected. To verify that these spectral bands are not artifacts or limited to very few spectra, we acquired hyperspectral images corresponding to the Amide-I region for all the IHC-identified cored plaques and their surrounding microenvironments (approximately 10,000 spectra per plaque region). To visualize the spatial distribution of parallel and antiparallel β-sheets, we calculated the ratios 1628:1660 and 1692:1660 respectively for each pixel, which have been shown to reflect the relative β-sheet populations in AD tissue^23, 24, 31^. The ratio images and corresponding IHC stained images for three representative plaques are shown in Figure 2. These three representative plaques are referred to as P_1_, P_2_, and P_3_ in the rest of the manuscript for clarity and are labeled in Figure 2 accordingly.

**Figure 1:**
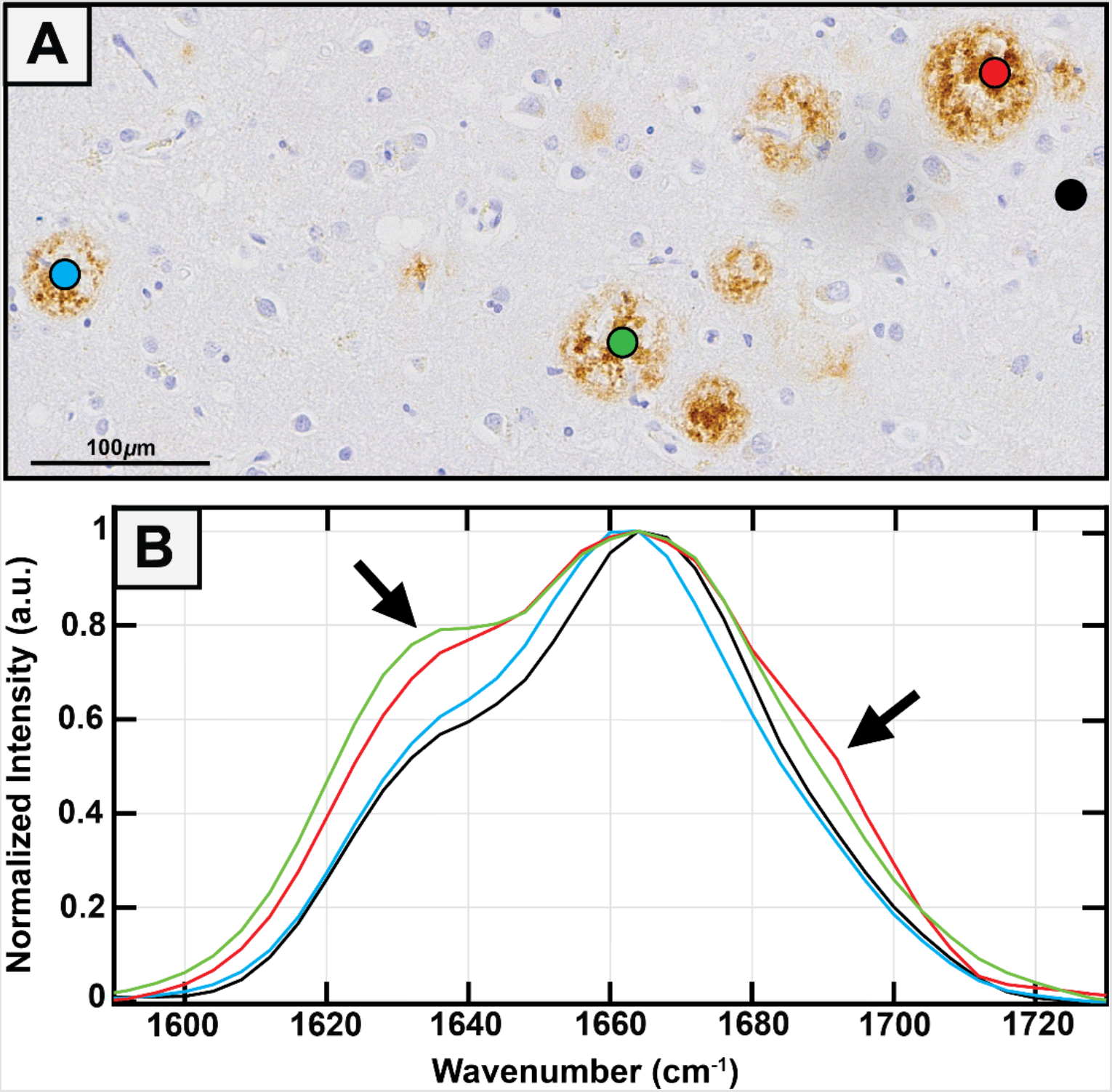
(A) Representative area of IHC stained plaques from frontal lobe. Three cored plaques are marked with blue (no β-sheet), green (β-sheet containing), and red (parallel and antiparallel β-sheet). For comparison, an outside area is marked with black. The scale bar is equal to 100 μm. (B) Average spectra from within each of the three plaque cores along with the outside area are stacked and shown.

**Figure 2:**
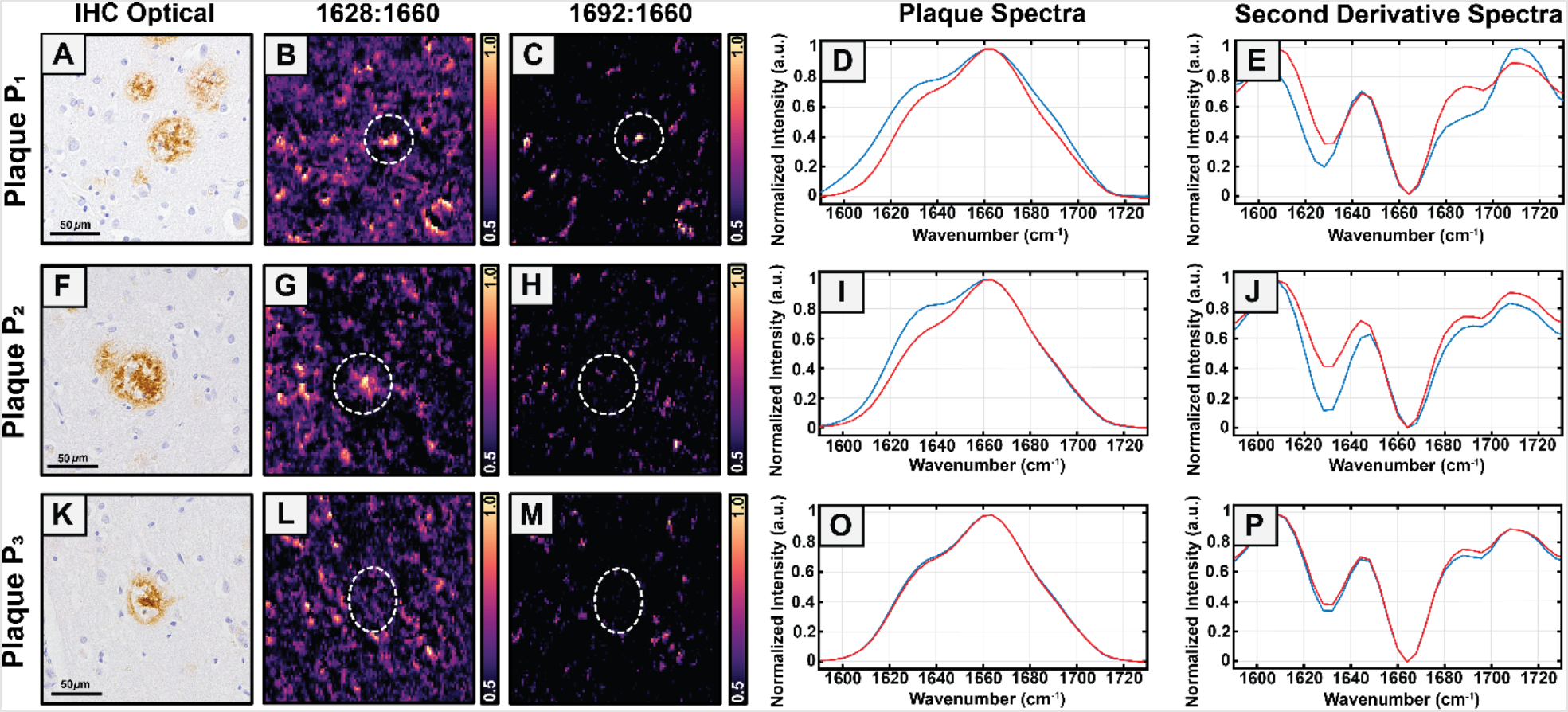
IHC stained images of three cored plaques (known as P_1_, P_2_, and P_3_) from the frontal lobe are shown (A, F, K). The scale bar is equal to 50 μm. Ratio images of 1628:1660 (β-sheet containing) (B, G, L) and 1692:1660 (antiparallel β-sheet containing) (C, H, M) are shown alongside the respective IHC images. Plaque P_1_ shows both parallel and antiparallel β-sheet signatures. Plaque P_2_ shows only parallel β-sheet signatures. Plaques P_3_ shows no significant β-sheet signatures. Average and second derivative spectra from plaques P_1_, P_2_, and P_3_ are also shown. The blue-line spectrum reflects the white, dashed circular plaque region and the red-line spectrum reflects the entire image field of view. Figure 2D, I, N compare the average spectra from within the plaque boundary to the average spectra of the field of view. Figure 2E, J, O compare the second derivative spectra from within the plaque boundary to the second derivative spectra of the field of view.

For each plaque we compare the average spectrum of pixels within the plaque boundary to the average spectrum of the entire area imaged, which is a 200 μm x 200 μm area around the plaque, including the plaque. For clarity, we refer to this as the microenvironment spectra for the rest of this article. Figure 2A-C show the IHC and ratio images corresponding to cored plaque P_1_ (Figure 2A-E), which clearly exhibits significant overall and antiparallel β-sheet signatures. Plaque P_2_ (Figure 2F-H) shows significant parallel β-sheet presence but lacks antiparallel intensity. Lastly, plaque P_3_ (Figure 2K-M) has no significant β-sheet presence in comparison to its microenvironment. This is also clearly reflected in the mean spectra and the second derivative spectra shown in Figure 2D-E, I-J, and N-O. Figure 2D-E show significant spectral differences in both the shoulder at ∼1628 cm^-1^ as well as at ∼1692 cm^-1^, which can be attributed to antiparallel β-sheet secondary structure. Figure 2I-J show significant differences at ∼1628 cm^-1^, but not at ∼1692 cm^-1^, and Figure 2N-O show no significant differences in terms of β-sheet between plaque average and microenvironment average. Additional IHC images, ratio images, spectra of plaques, and their second derivatives are shown in the Supporting Information (Figure S2). The cored plaques used for analysis can be categorized into one of the three subtypes (shown in Figure 2) based on the spectral variations observed. Namely, plaques that contain i. enhanced intensity at both ∼1630 cm^-1^ and ∼1692 cm^-1^, indicative of antiparallel β-sheets, ii. enhanced intensity only at ∼1630 cm^-1^ compared to their microenvironment, which we attribute to presence of overall β-sheets, and iii. no significant differences in spectra compared to the plaque microenvironment. It should be noted that the spectral ratios and the second derivatives plotted in Figure 2 are semi-quantitative metrics for assessing relative populations of protein secondary structure. The second derivative spectra of plaques P_1_, P_2_ and P_3_ from Figure 2 are plotted together in Figure S3 for comparison, which clearly highlights this spectral variation. The gold standard in IR spectroscopy for extracting secondary structure information is spectral deconvolution through band fitting^36^. To verify if the observed differences between plaques can indeed be attributed to a change in the spectral composition, we therefore employed spectral fitting to deconvolute the plaque and corresponding microenvironment mean spectra.

The spectra were fitted to three components, or sub-bands, which are consistent with observations from the experimental spectra and their second derivatives in Figures 1 and 2. The fit results for the three plaques (P_1_, P_2_, and P_3_) in Figure 2 are shown in Figure 3A-C. From the spectral fits, the contribution of β-sheets (overall and antiparallel) to the spectra was determined as the area under the curve (AUC) of the corresponding fitted band. The β-sheet populations thus determined for plaques P_1_, P_2_ and P_3_, are shown in Fig. 3D-E. We observe that plaque P_1_contains elevated populations of both antiparallel and overall β-sheets compared to its microenvironment, plaque P_2_ exhibits only elevated overall β-sheet content, while plaque P_3_ exhibits no significant differences from its microenvironment. This is in agreement with the conclusions drawn from the ratio images and plaque spectra.

**Figure 3:**
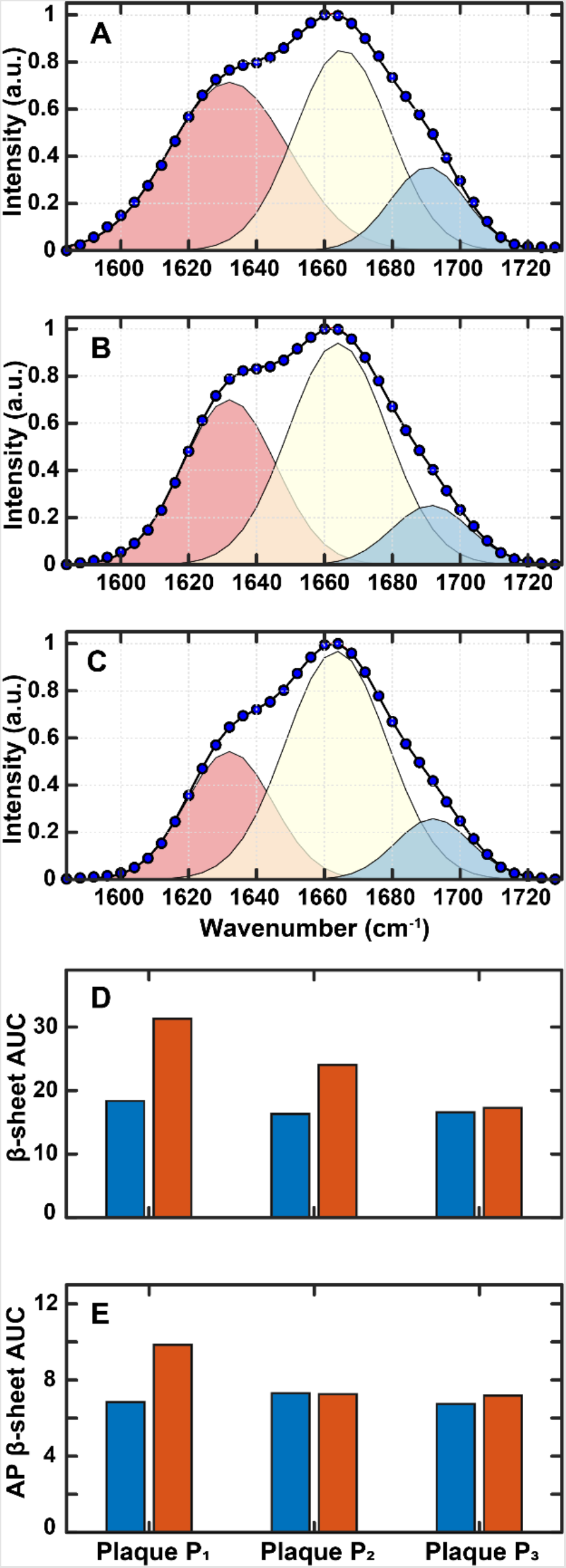
Spectral fit results from plaques P_1_, P_2_, and P_3_ are shown (A, B, and C) and labeled accordingly. Overall (D) and antiparallel (E) β-sheet population for plaques P_1_, P_2_, and P_3_ (red) and their corresponding microenvironments (blue).

We consequently expanded this analysis to all 80 plaques studied in this report. For a given plaque, we compared the β-sheet AUCs (overall and antiparallel) of the plaque core to its microenvironment and also to the mean AUC from all plaque microenvironments, and subsequently classified each plaque as β-sheet containing, antiparallel β-sheet containing, or β-sheet lacking. The results are shown in Figure 4. Details about the fitting approach are provided in the Supporting Information. Of the 80 plaques studied, 52 plaques show elevated overall β-sheet signatures, which is consistent with our previous observations^24^. Furthermore, out of these 52 plaques, 28 plaques also exhibit significant antiparallel β-sheet signatures. From the fitting analysis, we can infer that the spectral variations between the plaques can indeed be attributed to change in relative percentages of β-sheets. This validates the interpretation of the spectral variations in terms of changes in secondary structure. Taken together, the above analyses confirm that the average plaque spectra reflect any significant presence of β-sheets, overall and/or antiparallel, and can be used to determine the relative abundance of the same within a plaque. The deviations from the ideal parallel β-sheet structures of fibrils in plaques are not possible to determine from spatially averaged spectroscopy of in-vitro models, which underscores the significance of these results. We discuss the implications of these observations below.

**Figure 4:**
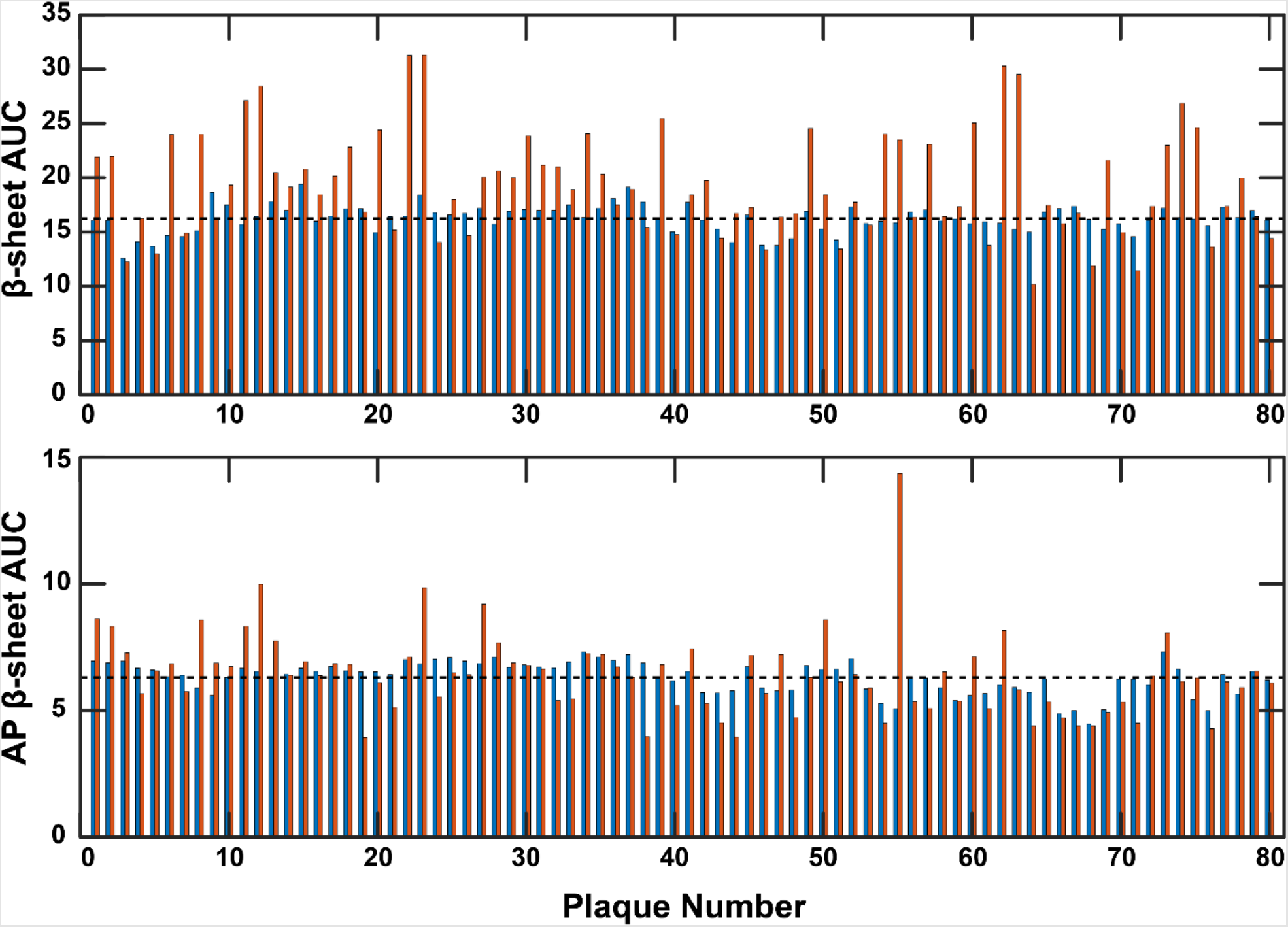
For all 80 plaques studied, a comparison of the β-sheet AUCs (overall and antiparallel) of the plaque core to its microenvironment and also to the mean AUC from all plaque microenvironments, and subsequently classified each plaque as β-sheet containing, antiparallel β-sheet containing, or β-sheet lacking.

The aggregation of amyloid proteins, specifically 40 or 42 residue wild-type Aβ peptides, has been extensively studied in-vitro with several techniques including ssNMR, cryo-EM, and IR spectroscopy^7, 10, 11, 30, 35^. It is believed that the aggregation proceeds from disordered monomers to β-sheet containing oligomers and then to fibrils, where the dominant structural motif is the parallel β-sheet^10, 33, 34, 37, 38^. Thus, observation of parallel β-sheet structure in plaques is expected and consistent with the known in-vitro structures of fibrils. It is also in agreement with prior IR imaging studies of AD tissues which have observed distinct parallel β-sheet character in plaques^23-25^. A lack of β-sheet peaks from plaques can arise from either the presence of more non-fibrillar deposits instead of fibrils or can be an inherent structural facet of the fibrils themselves. We have identified this heterogeneity in plaques in our recent work, specifically that individual fibrils, at early stages of aggregation, can often lack β-sheet structure^13^, leading to amyloid plaques exhibiting no enhanced β-sheet signature in IR spectra. The results shown here provide a validation of the generality of our previous findings. While 40 or 42 residue wild-type Aβ peptides form parallel β-sheets in-vitro, it is known that smaller fragments of Aβ spanning the amyloidogenic segment, such as Aβ 16-22 or 11-28, can form antiparallel β-sheets^39, 40^. These fragments have also been isolated from AD brain tissues^41^; hence observation of antiparallel structure can thus arise from the presence of these shorter fragments in plaques. However, the IHC antibody used in this work specifically binds to residues 1-4 in the Aβ sequence^26^; thus, it is unlikely that the plaques identified through IHC contain a large fraction of smaller Aβ fragments. The structure of oligomers in comparison to fibrils has been less studied, and different reports have described a distribution of structures for oligomers, including β-turns and antiparallel β-sheets^33, 38, 42^. We have not used any additional characterization of plaques besides staining to determine if they exclusively contain fibrils; therefore, it is possible that some oligomeric species are present in the plaques. It has been demonstrated that regions around the plaque core contain oligomeric Aβ, creating a halo or ring-like structure around the core^6, 43^. We observe this morphological feature in our IHC images (Figures 2A, F, K) but not in the IR ratio images (Figures 2B, G, L) which indicates that oligomeric species in tissues do not contain any significant amount of β-sheet, parallel or antiparallel. This is also consistent with prior reports on mouse models with FTIR imaging, where diffuse plaques, which contain mainly amorphous oligomeric/prefibrillar aggregates of Aβ, were found to exhibit no characteristic spectral features^22^. We therefore conclude that the antiparallel signatures observed herein are most likely arising out of fibrillar aggregates. The same argument can be extended for plaques that exhibit no distinct β-sheet signatures. Since oligomeric/non-fibrillar species primarily form a halo around the plaque, it is likely that the lack of β-sheet bands from the plaque core is a feature of fibrillar aggregates. In-vitro structural characterizations overwhelmingly show that antiparallel structure is rare in the longer isoforms of Aβ. These longer isoforms typically exhibit the parallel cross β-structure, which is dominant and contains higher structural order. However, there are reported exceptions such as those seen in recent work by Herzberg et. al^16^. They show the co-existence of parallel and antiparallel β-sheets within large amorphous aggregates at early stages of aggregation, which also contain fibrils. Ruggeri et. al. have also investigated amyloid aggregation with nanoscale IR spectroscopy and have demonstrated that even after the structural conversion from oligomers to fibrils, the latter contain antiparallel signatures^14^. Fibrils from racemic mixtures of Aβ peptides have been recently shown to contain antiparallel structure^44^. Irizarry et. al. have shown that fibrils seeded from cerebral vascular tissues of CAA patients, another hallmark of AD, contain a mixture of parallel and antiparallel β-sheet structures^18^. In fact, the Iowa mutant of Aβ, also implicated in CAA, has been shown to form transient fibrils with antiparallel structure that eventually convert to parallel β-sheets^19^. CAA pertains to vascular amyloids deposits; however, the presence of antiparallel structure in CAA-related amyloid variants nonetheless points to the possibilities that these structures can also exist for wild-type Aβ in plaques. Recent cryo-EM investigation of fibrils seeded from AD brain extracts shows evidence of intramolecular β-hairpin conformation in the outer cross-β layers, leading to a fibril structure that contains antiparallel structural elements^17^. To the best of our knowledge, this work is the first demonstration of significant prevalence of antiparallel β-sheets in ex-vivo specimens, specifically in fibrillar amyloid plaques. It has been suggested that wild-type Aβ fibrils with antiparallel structure may exist but could also be transient in nature. It is also possible that the relative population of antiparallel fibrils is significantly smaller than more stable parallel aggregates, and as a result they are not detected in conventional spatially averaged measurements. It should be noted in this context that amyloid aggregation involves a dynamic equilibrium between several aggregate species; the capability of spatially resolving this structural ensemble allows for isolating spectral signatures of transient and/or less prevalent species. However, that does not necessarily guarantee that presence of such species aggregation ensembles in-vitro will be identifiable by IR imaging, since these measurements do not probe structures of isolated/individual aggregates. Therefore, to test the hypothesis that deviations from parallel fibril structures exist in wild-type Aβ and verify that such deviations can be identified using IR imaging, we expanded our analysis to Aβ 1-42 fibrils aggregated in-vitro.

The aggregation was formed at a concentration of 100 μM in 2 mM HCl (pH 2.0); fibrils formed at this pH have been suggested to be morphologically similar to those isolated from AD brain^45^. Fibrils were observed after ∼3 hours of aggregation, and their presence in the aggregation mixture was verified by Atomic Force Microscopy (AFM). A representative AFM topograph of fibrils is shown in Figure 5A. To test if these fibrillar specimens could potentially contain antiparallel structure that might be captured by IR imaging, we acquired spatially resolved IR spectra from 3 different spatial locations. For each spatial location, 100 spectra were acquired spanning an area of 10 μm x 10 μm, approximately the dimensions of plaque cores. The mean spectra of each location (Figure 5B-D) indicate that there can be significant spectral variations between different locations within the ensemble. To validate if these variations can be attributed to relative changes in antiparallel β-structure, we fit the mean spectra using the same procedure used for plaques (vide supra). The fit results, shown in Figure 5B-D, demonstrate that the fibril spectra can be decomposed into a peak at the typical parallel β-sheet wavenumber of ∼1630 cm^-1^ and another at ∼1660 cm^-1^ corresponding to random coils and β-turns, similar to the spectra in AD tissues. The overall and antiparallel β-sheet populations determined for areas 1, 2, and 3 (vide supra) are shown in Figure 5E. Interestingly, all the spectra exhibit an additional peak at ∼1692 cm^-1^, which varies significantly in intensity between the different spatial locations/aggregates. This observation indicates that even in wild-type Aβ fibrillar aggregates, a fraction of the ensemble can contain antiparallel structure and is in agreement with our observations in ex-vivo tissue sections. It must be noted that the above measurements are not aimed at providing definitive statistics on the specific fraction of fibrils that contain antiparallel structure, but to merely identify the presence of antiparallel structure in fibrils. Furthermore, the fibrils studied here are in early stages of maturation and can undergo additional evolution that alters the relative population of antiparallel structure. We have recently demonstrated such structural evolution in the fibrillar phase for Aβ 16-22^40^. Nonetheless, these results demonstrate deviation from the expected parallel structure in wild-type Aβ 1-42 fibrils. Amyloid aggregates in AD plaques can exist in a range of aggregation states, from early stage to mature fibrils. It is therefore likely that fibrils with antiparallel character, such as those studied here, will be found in some plaques, and these results hence offer a validation and/or rationalization for identification of antiparallel structure in plaques with spatially resolved IR spectroscopy. We aim to study the exact prevalence of antiparallel character in Aβ fibrils and their evolution in more detail in future work.

**Figure 5:**
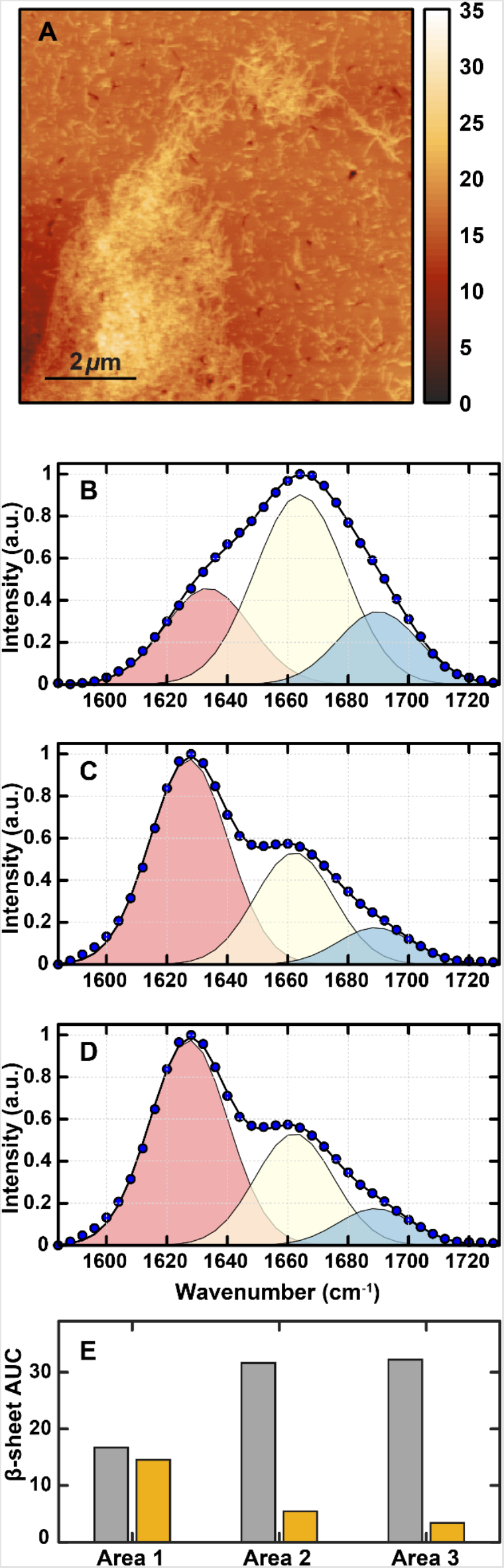
(A) AFM topograph of Aβ 1-42 fibrils. The scale bar is equal to 2 μm. (B-D) Spectral fit results of mean IR spectra acquired from 3 different spatial locations. (E) Overall (gray) and antiparallel (gold) β-sheet population for areas 1, 2, and 3.

Taken together, the results reported herein provide a link between the structure of in-vitro amyloid aggregates and those found in AD brain. The presence of off-pathway species as well as intermediate structures have been identified during amyloid aggregation in-vitro; however, the presence of such species in AD plaques has been rarely explored. Therefore, the structural distribution of amyloid aggregates in plaques is poorly understood, regardless of being imperative for the development of therapeutic interventions of AD that specifically target amyloid plaques. The study presented here precisely addresses this gap in knowledge, and demonstrates that antiparallel β-sheets, which are believed to be largely transient/intermediate species during amyloid aggregation and have been shown to be equally neurotoxic as aggregates with parallel β-structure, can exist in cored plaques. Our findings highlight the possibility of chemically subtyping plaques, wherein the chemistry and/or the secondary structure of amyloid aggregates in plaques is the key categorizing metric instead of their morphologies in histochemical stains. The correlation of plaques of a certain chemistry in different stages or types of AD can potentially unlock novel therapeutic strategies for AD and associated dementia. This work also paves the way to test the correlation of plaque secondary structure with varying stages of AD pathogenesis in future studies.

## Materials and Methods

### AD Tissue Samples

Formalin fixed paraffin embedded (FFPE) human frontal lobe diseased tissue samples were purchased from Advanced Tissue Services (Tucson, AZ). One frontal lobe sample is from an 89-year old female AD patient; the other frontal lobe sample is from an 81-year old male AD patient. The postmortem tissue specimens studied in this report were deidentified and were determined not to be human subjects research by the Office of Research Compliance at the University of Alabama. The tissues were deparaffinized in n-hexane for 24 h before storage under a mild vacuum prior to IR imaging. For IHC staining, sections adjacent to those used for IR measurements were stained with the anti-amyloid MOAB-2 antibody (Sigma-Aldrich) at the University of Alabama at Birmingham pathology core research lab. The IHC stained sections were used as a visual guide for identifying Aβ plaque location and morphological type. Brightfield images of IHC stained tissues were acquired at 40× magnification with an Olympus BX43 microscope using manualWSI software (Microvisioneer) for image stitching.

### IR Imaging

IR images were acquired using a home-built stage scanning IR microscope. The microscope design is based upon work published by Yeh and co-workers^32, 46^. The microscope uses a quantum cascade laser system (LaserTune, Block Engineering) that is tunable from ∼1000 to 1800 cm^−1^ for illumination, and a thermoelectrically (TE) cooled mercury cadmium telluride (MCT) detector (Vigo Photonics) and is equipped with a 0.71NA objective. The hyperspectral data sets were acquired within the spectral range of 1584−1730 cm^−1^ at 4 cm^-1^ spectral and 2 μm spatial resolution. All measurements were made in transflection mode on IR reflective low-emissivity slides (mirrIR, Kevley Technologies).

### Data Processing

All fitting procedures, image generation, and analyses were performed using MATLAB software. Ratio images were created by ratioing peaks of interest within the Amide-I region that correspond to relevant signatures of known protein secondary structures. These ratio images provide a method for normalization between the local areas of analysis and the entire tissue to account for variations in density and thickness within the entire sample. The following intensity ratios were used: 1628:1660 and 1692:1660. These ratios represent signatures corresponding to parallel and antiparallel β-sheets, respectively. Spectra were smoothed using a 5-point moving average filter and a linear baseline correction was performed.

### Aβ42 aggregation

Aβ42 (rPeptide, USA) was first treated with 1,1,1,3,3,3-hexafluoroisopropanol (HFIP) for 15 minutes at room temperature. Then HFIP was evaporated by keeping the vial in a vacuum desiccator. After complete removal of HFIP, a stock solution of 1 mM Aβ42 in 10 mM HCl was prepared and incubated at 37°C without any agitation for aggregation.

### Sample preparation for AFM-IR experiment

Samples were prepared on ultra-flat gold substrate (Platypus Technologies, USA) by depositing 10 μL of 100 μM aggregate solution and incubated for 5 minutes. The samples were rinsed with 100 μL of milli-q water and dried under nitrogen stream before imaging.

## Acknowledgment

This work was supported by the National Institutes of Health (Award 1 R35 GM138162 to A. G.). The authors thank Dr. Dezhi Wang at UAB Pathology Core Facility for help with IHC staining.

## Supporting Information

IHC optical image and ratio images for full tissue sample sections; additional IHC optical images, 1628:1660 ratio images, 1692:1660 ratio images, mean spectra, and second derivative spectra for plaques exhibiting antiparallel β-sheet signatures, parallel β-sheet only, and no β-sheet; additional details regarding deconvolution of plaque spectra using Gaussian band fitting; stacked second derivative spectra from Figure 2.

## Data Availability

All data/results needed to evaluate the conclusions drawn in this communication are included in the manuscript and the Supplementary Information file. The datasets and any other additional data are available from the corresponding author upon reasonable request.

