## Supporting Information for "Intermediate antiparallel beta structure in amyloid plaques revealed by infrared spectroscopic imaging"

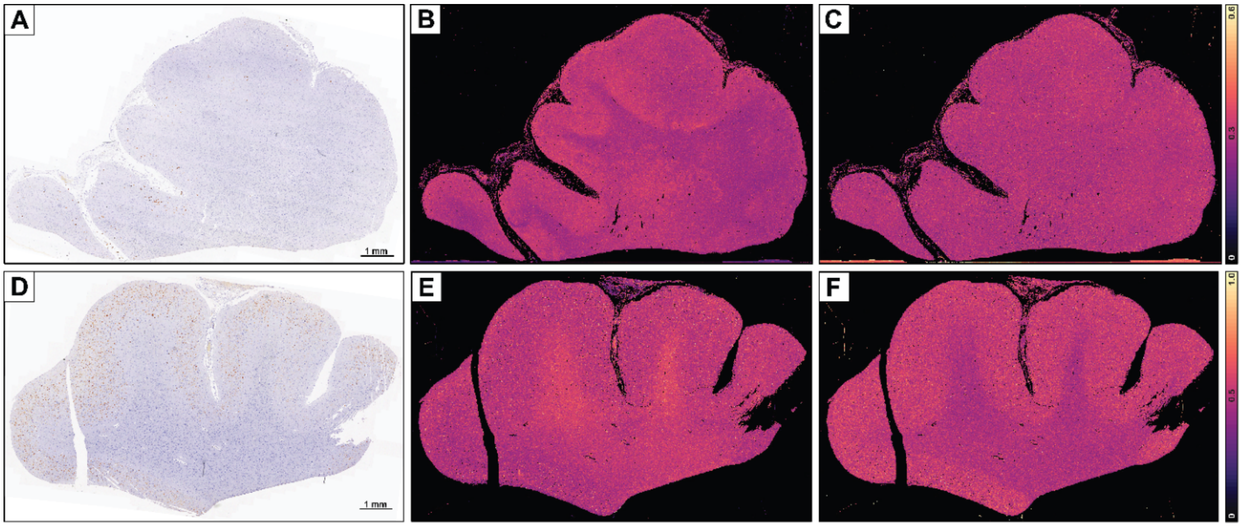


**Figure S1:** IHC stained full tissue optical images of the two frontal lobe sections used for this analysis are shown (A,D). The scale bar is equal to 1 mm. Ratio images of 1628:1660 $cm^{-1}$ (overall $\beta$-sheet) (B, E) and 1692:1660 $cm^{-1}$ (overall and antiparallel $\beta$-sheet) (C, F).

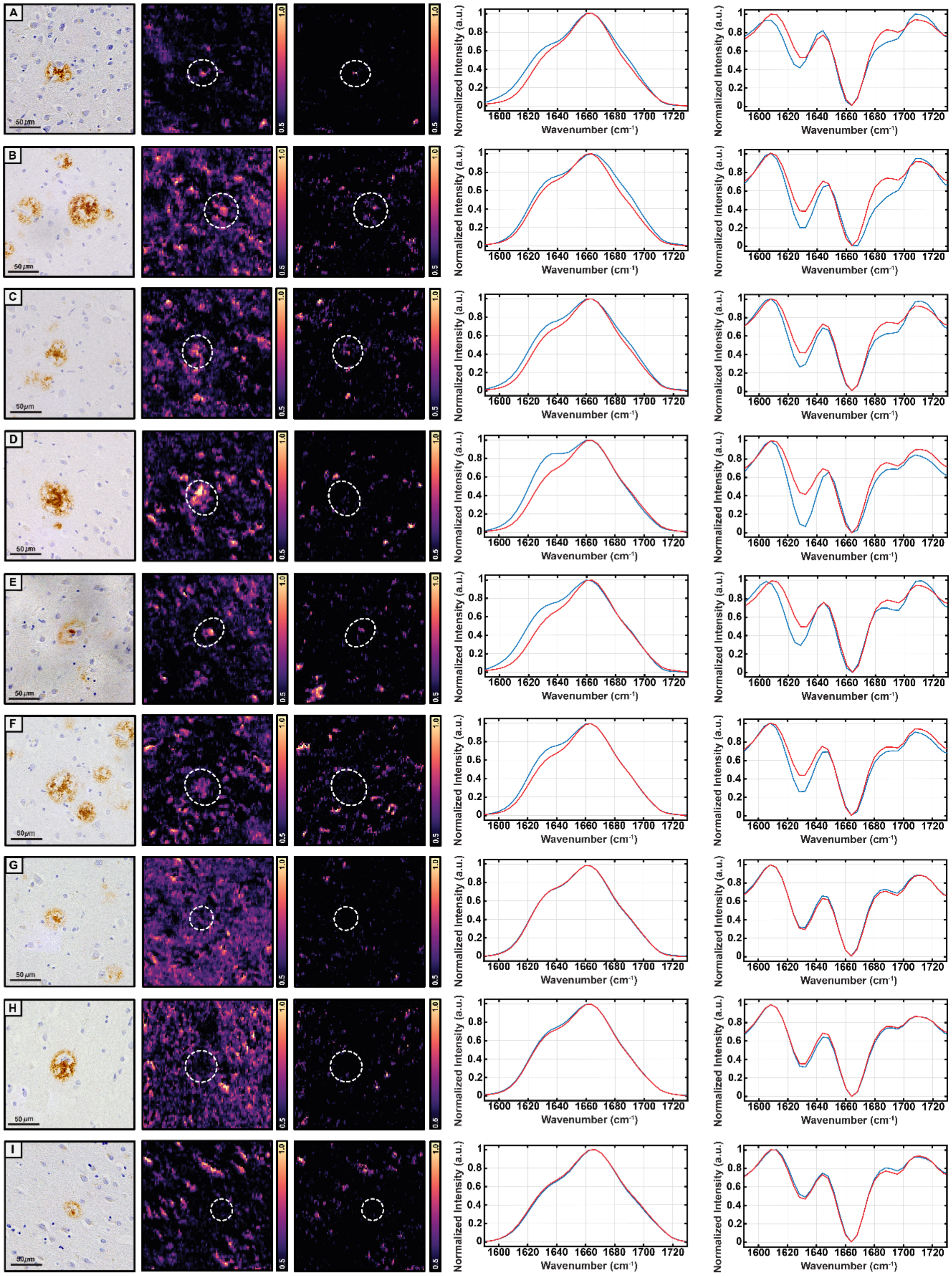


**Figure S2**: IHC stained images and ratio images of additional cored plaques from frontal lobe are shown. Ratio images of 1628:1660 cm$^{-1}$ (β-sheet) and 1692:1660 cm$^{-1}$ (antiparallel β-sheet) are shown alongside their respective plaques. Average and second derivative spectra are also shown.

**Deconvolution of plaque spectra using Gaussian band fitting:**

For quantitative assessment of the relative β-sheet content of the plaques, the mean spectra from the plaque core and their microenvironments were fitted to a sum of three Gaussian bands:

The mean spectra shown in Figures 1-2 in the main manuscript were fit to a sum of four constituent Gaussian peaks:

$$S = \sum_{n=1}^{3} a_n e^{-\left(\frac{w-w_{0,n}}{c}\right)^2}$$

where $a_n$ is the amplitude of the n-th peak, and $w_{0,n}$ the corresponding center wavenumber. The spectra were normalized to the maximum intensity of the Amide-I band prior to fitting to mitigate any intensity fluctuations from tissue thickness variations. The peak positions identified in the second derivative spectra, shown in Figure 2 in the manuscript, were used as a starting point for the fitting procedure. The area under the curve (AUC) of each fitted band represents the concentration/abundance of the corresponding structural moeity. Hence, to determine if a plaque is β-sheet depleted or has enhanced β-sheet content, we compared the AUCs of the bands centered at ~1630 $cm^{-1}$ and ~1690 $cm^{-1}$ between the plaque and a 200 μm x 200 μm area around the plaque (which also included the plaque). For clarity, we refer to this as the plaque microenvironment. For a plaque to be counted as β-sheet 'rich', it had to exhibit larger AUC of the ~1630 $cm^{-1}$ band than its microenvironment, and also had to have an AUC higher than the mean AUC value obtained for all the microenvironments. A similar metric was used to categorize plaques as antiparallel β-sheet containing, except that the AUC of the ~1690 $cm^{-1}$ band was used. The mean spectra from different in-vitro Aβ fibrillar aggregates were fitted using an identical procedure.

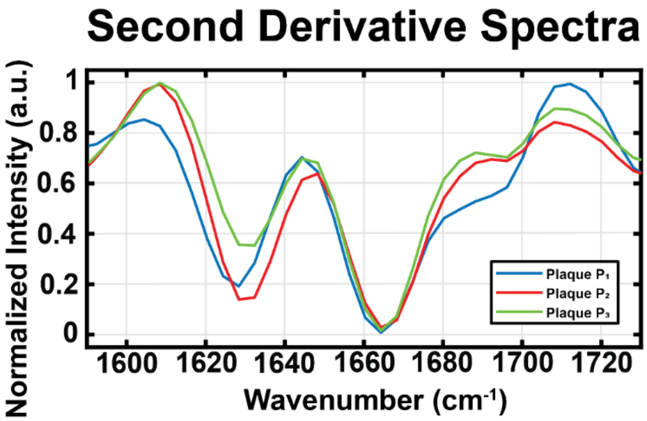


**Figure S3**: Second derivative spectra from plaques $P_1$, $P_2$, and $P_3$ are stacked and shown for direct comparison.

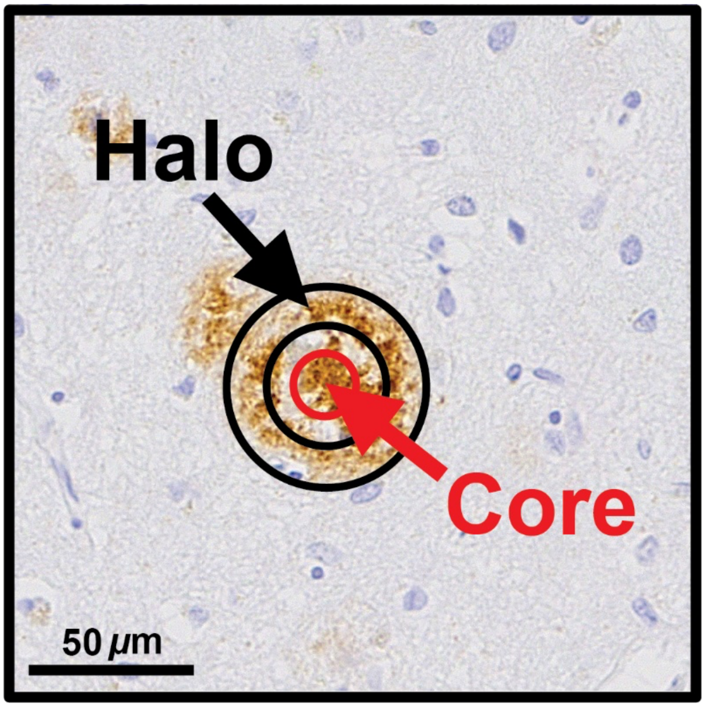


**Figure S4**: Plaque P2 is shown and annotated to better elaborate on the morphological features of a cored plaque. The plaque "core" is marked and shown to be within the red circle. The core is referring to the central, densest area of the plaque. The plaque "halo" is marked and shown to be within the two black circles. The halo is referring to the ring-like feature surrounding the core of the plaque.
